# In vitro: Natural Compounds (Thymol, Carvacrol, Hesperidine, And Thymoquinone) Against Sars-Cov2 Strain Isolated From Egyptian Patients

**DOI:** 10.1101/2020.11.07.367649

**Authors:** M.G Seadawy, A.F Gad, M.F Elhoseny, B.El ELharty, M.D Shamel, Abdo A. Elfiky, Aya Ahmed, Abdel Rahman N. Zekri

**Affiliations:** Main chemical laboratories, Egypt Army; Biophysics Department, Faculty of Science, Cairo University; Molecular Virology and Immunology Unit, Cancer Biology Department, National Cancer Institute, Cairo University, Cairo, Egypt; National Cancer Institute, Cairo University, Giza, Egypt

**Keywords:** COVID-19, SARS-CoV-2, Natural products, antivirals, docking, plaque reduction assay

## Abstract

The current pandemic of the coronavirus disease-2019 (COVID-19) has badly affected our life during the year 2020. SARS-CoV-2 is the primary causative agent of the newly emerged pandemic. Natural flavonoids, Terpenoid and Thymoquinone are tested against different viral and host-cell protein targets. These natural compounds have a good history in treating Hepatitis C Virus (HCV) and Human Immunodeficiency Virus (HIV). Molecular docking combined with cytotoxicity and plaque reduction assay is used to test the natural compounds against different viral (Spike, RdRp, and M^pro^) and host-cell (TMPRSS II, keap 1, and ACE2) targets. The results demonstrate the binding possibility of the natural compounds (Thymol, Carvacrol, Hesperidine, and Thymoquinone) to the viral main protease (M^pro^). Some of these natural compounds were approved to start clinical trail from Egypt Center for Research and Regenerative Medicine ECRRM IRB (Certificate No.IRB00012517)

## Introduction

By the end of 2019, an outbreak of a novel coronavirus (SARS-CoV-2) in Wuhan city in China was detected and spread all over the world ^1^. On 10^th^ October 2020, the number of confirmed cases of coronavirus disease (COVID-19) reached more than 37 M worldwide with +1M total death, as reported in the World Health Organization (WHO). The associated pneumonia with the novel viral infection, COVID-19, is divided into three phases that correspond to different clinical stages of the disease ^2^. Stage 1 is the asymptomatic stage, where the inhaled virus binds to nasal epithelial cells in the nasal cavity and starts replicating. Stage 2 is the upper airway stage, where the virus propagates, migrates down the respiratory tract along the conducting airways, and a more robust innate immune response is triggered. About 20% of the infected patients will progress to stage 3 disease and develop pulmonary infiltrates. Some of these patients will develop a very severe disease as the virus reaches alveoli in the lung and infects alveolar type II cells in peripheral and sub-pleural areas of the lung ^3^. SARS-CoV-2 propagates within type II cells, large numbers of viral particles are released, and the cells undergo apoptosis and die. Therefore, the spectrum of symptomatic COVID-19 ranges from mild respiratory tract infection to severe pneumonia that may progress to fatal respiratory syndrome and multi-organ malfunctions ^2^.

Thymol, known as 2-isopropyl-5-methylphenol, is a natural monoterpenoid phenol derivative of Cymene. It is found in thyme oil and extracted from Thymus vulgaris ^4^. Thymol is a white crystalline substance that has a pleasant aromatic odor. Thymol also provides the distinctive and robust flavor of the culinary herb thyme.

Carvacrol is known as monoterpenoid phenol and extracted from Oregano. It has a characteristic pungent and warm odor ^5^.

Hesperidine is a common flavone glycoside found in citrus fruit such as lemons and sweet oranges ^6,7^. It has several pharmacological activities such as antihyperlipidemic, anti-atherogenic, venotonic, antidiabetic, cardioprotective, anti-antihypertensive, and inflammatory actions ^6,7^. The anti-inflammatory activity of hesperidin was mainly attributed to its antioxidant defense mechanism and suppression of pro-inflammatory cytokine production ^6^. Hesperidin exhibited antiviral activity against the influenza virus through a significant reduction of viral replication.

Nigella sativa (NS) contains many active molecules, such as thymoquinone (TQ), two forms of alkaloids: isoquinoline alkaloid that includes nigellicimine, nigellicimine n-oxide and pyrazol alkaloid that includes nigellidine and nigellicine ^8,9^. TQ is the most abundant constituent in the volatile oil of *Nigella sativa* seeds, and most of the herb’s properties are attributed to it ^10,11^. It has been reported that NS oil can decrease the viral count of HCV in patients received capsules of NS oil (450 mg) three times a day over a 3-month period ^12^. Moreover, two clinical studies documented to sustained sero-reversion of the HIV virus over treatment period of 6 to 12 months ^13–15^.

Molecular docking represents a promising *in silico* method used to predict the binding affinities of small molecules to proteins as a first step in structure based drug design ^16–21^. This study investigated many active ingredients that showed antiviral activities against SARS-CoV-2, such as Thymol, Carvacrol, Hesperidine, and Thymoquinone. Molecular docking is used to test the binding affinities of these natural product derived compounds against different viral and host cell proteins. Additionally cytotoxicity assay and plaque reduction assay are used to verify their antiviral activity against SARS-CoV-2 collected from Egyptian patients.

## Materials and methods

### In silico testing

Before performing the docking studies, the tested compounds are retrieved from the PubChem database and then prepared using PyMOL software ^22,23^. The structures of the proteins are downloaded from the protein data bank ^24^. Autodock Tools are used to prepare the docking input files after adding charges ^25^. The docking experiments are performed utilizing Autodock Vina software in triplicates to have a clear depiction of the mode of action of compounds against their protein targets ^26^. SARS-CoV-2 Spike protein (S), The RNA dependent RNA polymerase (RdRp), the main protease (M^pro^) are used as protein targets due to its critical role in maintaining the viral infection ^27–31^. Additionally, the host-cell receptors, Transmembrane protease, serine 2 (TMPRSS II), Kelch-like ECH-associated protein 1 (KEAP 1) and the Angiotensin Converting Enzyme 2 (ACE2) are targeted due to its fundamental in viral recognition and maintain infectivity for SARS-CoV-2 ^32–35^. For each compound, ten interactions were generated and one with best binding affinity was selected. PyMOL software was used to represent and analyze the docking complexes.

### Experimental section

All the chemical compounds are purchased from different sources as follow; Thymol purchased from upnature as THYME 100% pure and natural 118 ml, Carvacrol purchased from Zane Hellas as oil of Oregano 30 ml, Hesperidin purchased from science-based nutrition as hesperidin methyl chalcone 500 mg −60 veggie caps, and Thymoquinone purchased from prime natural black seed USDA organic 240 ml.

### A) Cytotoxicity assay

Samples were diluted with Dulbecco’s Modified Eagle’s Medium (DMEM). Stock solutions of the test compounds were prepared in 10 % DMSO in ddH_2_O. The cytotoxic activity of the extracts were tested in Vero E6 cells by using the 3-(4, 5-dimethylthiazol -2-yl)-2, 5-diphenyltetrazolium bromide (MTT) method ^36^ with minor modification. Briefly, the cells were seeded in 96 well-plates (100 µl/well at a density of 3×105 cells/ml) and incubated for 24 hrs at 37°C in 5%CO_2_. After 24 hrs, cells were treated with various concentrations of the tested compounds in triplicates. After further 24 hrs, the supernatant was discarded and cell monolayers were washed with sterile phosphate buffer saline (PBS) 3 times and MTT solution (20 µl of 5 mg/ml stock solution) was added to each well and incubated at 37°C for 4 hrs followed by medium aspiration. In each well, the formed formazan crystals were dissolved with 200 µl of acidified isopropanol (0.04 M HCl in absolute isopropanol = 0.073 ml HCL in 50 ml isopropanol). Absorbance of formazan solutions were measured at λ_max_ 540 nm with 620 nm as a reference wavelength using a multi-well plate reader. The percentage of cytotoxicity compared to the untreated cells was determined with the following equation.

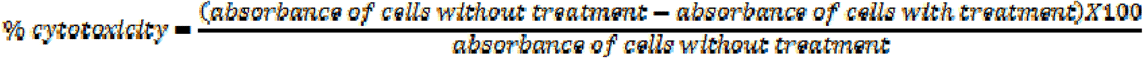

The plot of % cytotoxicity versus sample concentration was used to calculate the concentration which exhibited 50% cytotoxicity (IC50).

### B) Plaque reduction assay

Assay was carried out according to the method of ^37^ in a six well plate where Vero E6 cells (10^5^ cells / ml) were cultivated for 24 hrs at 37°C. Sever Acute Respiratory Syndrome Coronavirus (SARS-CoV2) virus was diluted to give 103 PFU /well and mixed with the safe concentration of the tested compounds and incubated for 1 hour at 37°C before being added to the cells. Growth medium was removed from the cell culture plates and the cells were inoculated with (100 µl /well) virus with the tested compounds, After 1 hour contact time for virus adsorption, 3 ml of DMEM supplemented with 2% agarose and the tested compounds was added onto the cell monolayer, plates were left to solidify and incubated at 37°C till formation of viral plaques (3 to 4 days). Formalin (10%) was added for two hours then plates were stained with 0.1 %crystal violet in distilled water. Control wells were included where untreated virus was incubated with Vero E6 cells and finally plaques were counted and percentage reduction in plaques formation in comparison to control wells was recorded as following:

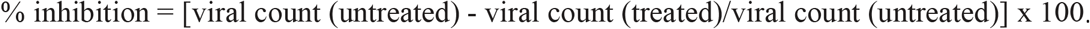

### Statistical analysis

Analysis was performed using Graphpad Prism 8.0.2. Data are represented as mean ± SD and statistical significance was evaluated using one-way ANOVA followed by tukey multiple comparison tests.

## Results and discussion

Many reports showed that different natural product derived compounds have promising results against inflammation and viral infections including the newly emerged viral infection (SARS-CoV-2) ^15,38–40^. Different natural compounds and their derivatives proved its binding affinity against different SARS-CoV-2 and host-cell targets ^38,39,41^.

Figure 1 shows the 2D structures of the selected natural compounds used in the current study against viral and host-cell protein targets.

**Figure 1.**
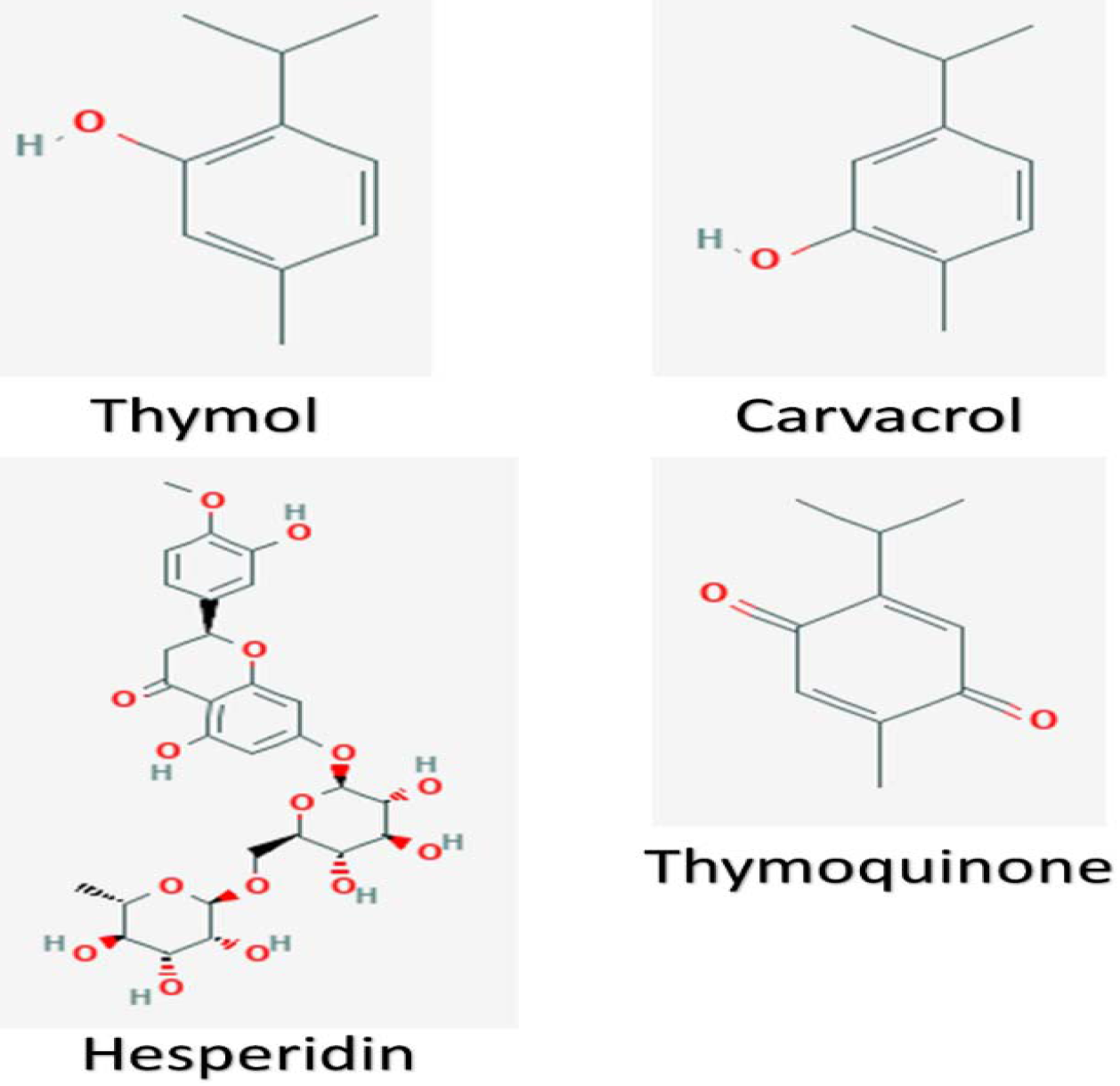
The structures of the natural compounds used in the study retrieved from the PubChem database.

Three SARS-CoV-2 proteins are targeted in this study including the viral, host-cell recognizing critical element, spike protein (Figure 2A), the vital viral enzyme RNA dependent RNA polymerase (RdRp), responsible for the polymerization of the complement RNA copy, and the Main protease (M^pro^) of SARS-CoV-2, which is critical for polyprotein processing (Figure 2B) ^42–47^. The spike protein trimers over the virions take different conformations during infection such as the prefusion and postfusion. The prefusion has open and closed conformations in which one or two of the receptor binding domains (RBD) are exposed (open) or immersed into the trimer (close) ^30,48^.

**Figure 2.**
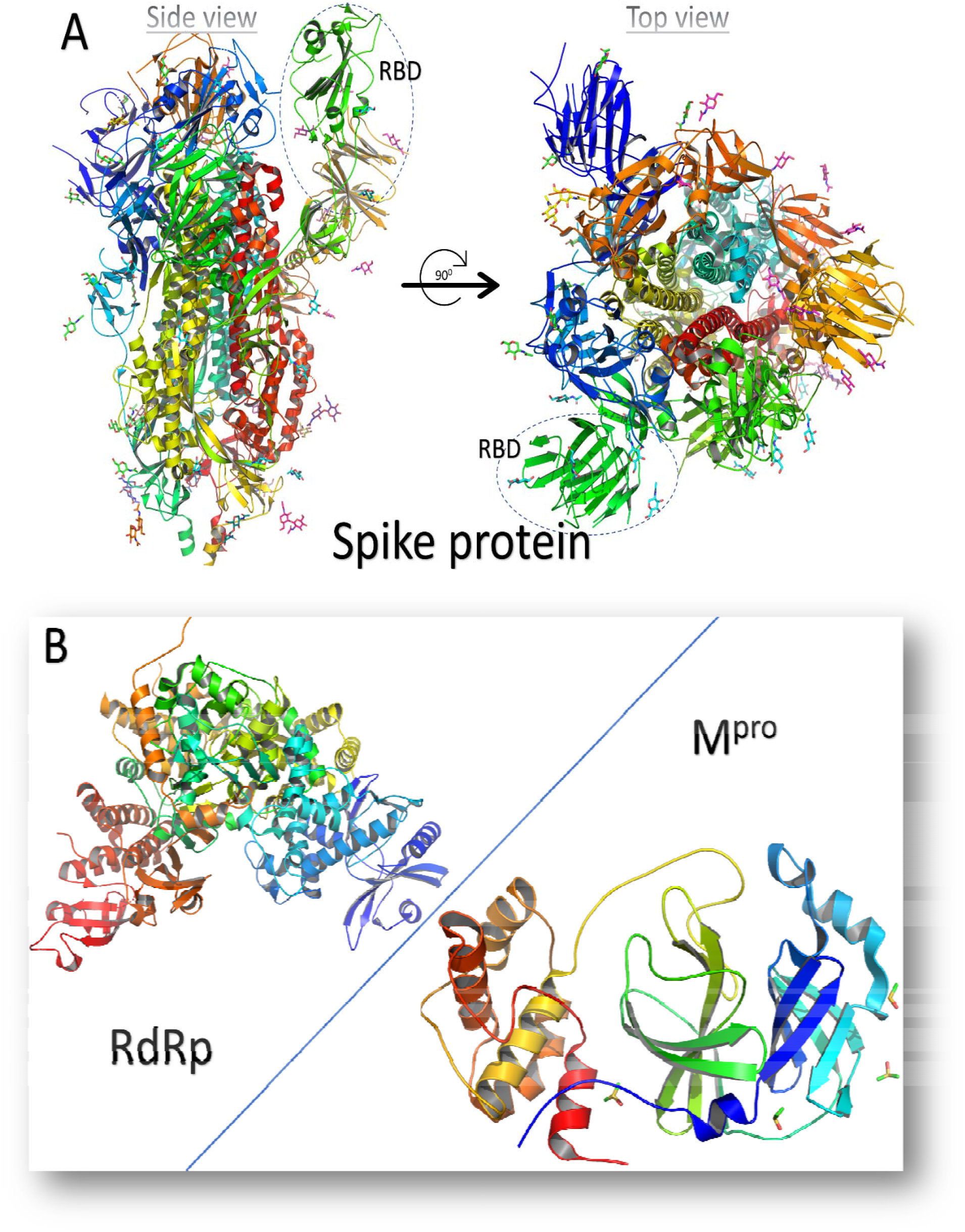
The 3D structures of the SARS-CoV-2 protein targets used in the study. **A)** The structure of SARS-CoV-2 Spike protein (PDB ID: 6VYB) represented by colored cartoons in side (left) and top (right) views. The receptor binding domain (RBD) is encircled in both views of the spike. **B)** The structure of the SARS-CoV-2 RdRp (PDB ID: 7BTF) and Mpro (PDB ID: 6Y84) depicted in colored cartoons. The structures are represented using PyMOL software.

Table 1 shows the binding affinity calculated using AutoDock Vina software for the docking of the natural compounds (Thymol, Carvacrol, Hesperidine, and Thymoquinone) against the SARS-CoV-2 M^pro^ as a protein target. The standard compound Chloroquine is used to assess the binding affinity of the natural compounds against the M^pro^. As reflected from the values, Carvacrol, Hesperidine, and Thymoquinone show comparable binding affinities (−7.0, −6.9, and - 6.9 kcal/mol, respectively) to SARS-CoV-2 M^pro^ compared to that of the standard compound (−7.2 kcal/mol). Thymol show slightly higher (worse) binding affinity value (−5.8 kcal/mol) compared to Chloroquine but still able to bind the SARS-CoV-2 M^pro^ tightly. Figures 3A and 3B show the 3D poses for the docking complexes. The four compounds (Thymol, Carvacrol, Hesperidine, and Thymoquinone) are able to bind to the active site of the M^pro^ (His41 and Cys145).

**Table 1.** The binding affinity (in kcal/mol) of the natural compounds against the main protease of SARS-CoV-2 calculated using AutoDock Vina software.

| Compound | Binding affinity(kcal/mol) |
| --- | --- |
| *Thymol* | -5.8 |
| *Carvacrol* | -7.0 |
| *Hesperidine* | -6.9 |
| *Thymoquinone* | -6.9 |
| *Chloroquine(standard)* | -7.2 |

**Figure 3.**
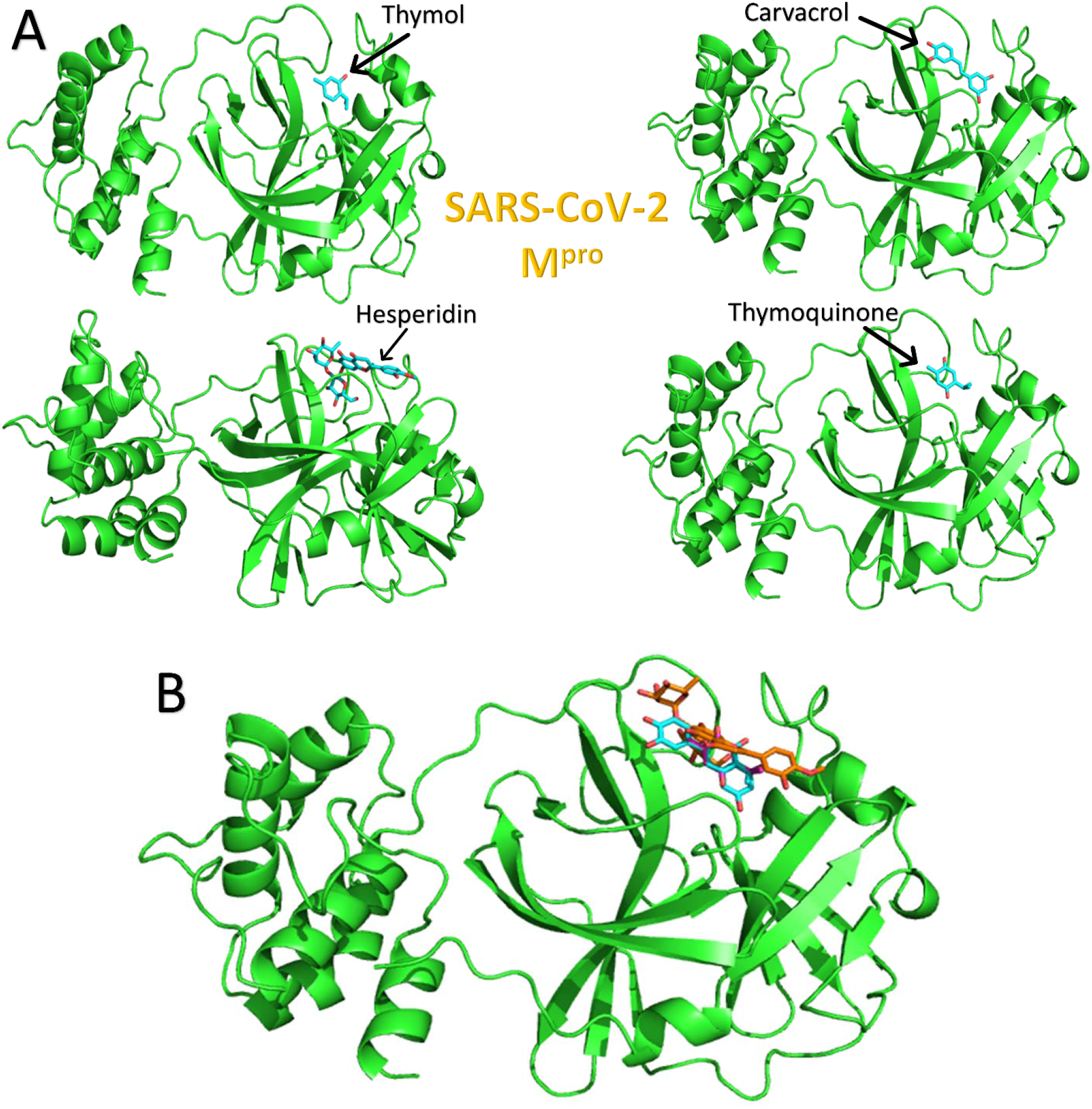
(A) The docking complexes formed after the docking of Thymol, Carvacrol, Hesperidine, and Thymoquinone into the SARS-CoV-2 M^pro^ active site. (B) The superposition of all the compounds. The protein is represented with green cartoon, while ligands are in cyan and orange sticks.

### The half maximal inhibitory concentration (IC50) of Thymol, Carvacrol, Hesperidin, and Thymoquinone

The effect of different concentrations of the compounds on the cellular proliferation of Vero E6 cell line following 24 h of treatment was determined using MTT assay.

Results of the present work revealed a concentration-dependent cytotoxic effect of Thymol (617 µM), Carvacrol (464 µM), Hesperidine (5.58 µM) and Thymoquinone (3.9 µM) on Vero E6 cell line as shown in figure 4.

**Figure 4:**
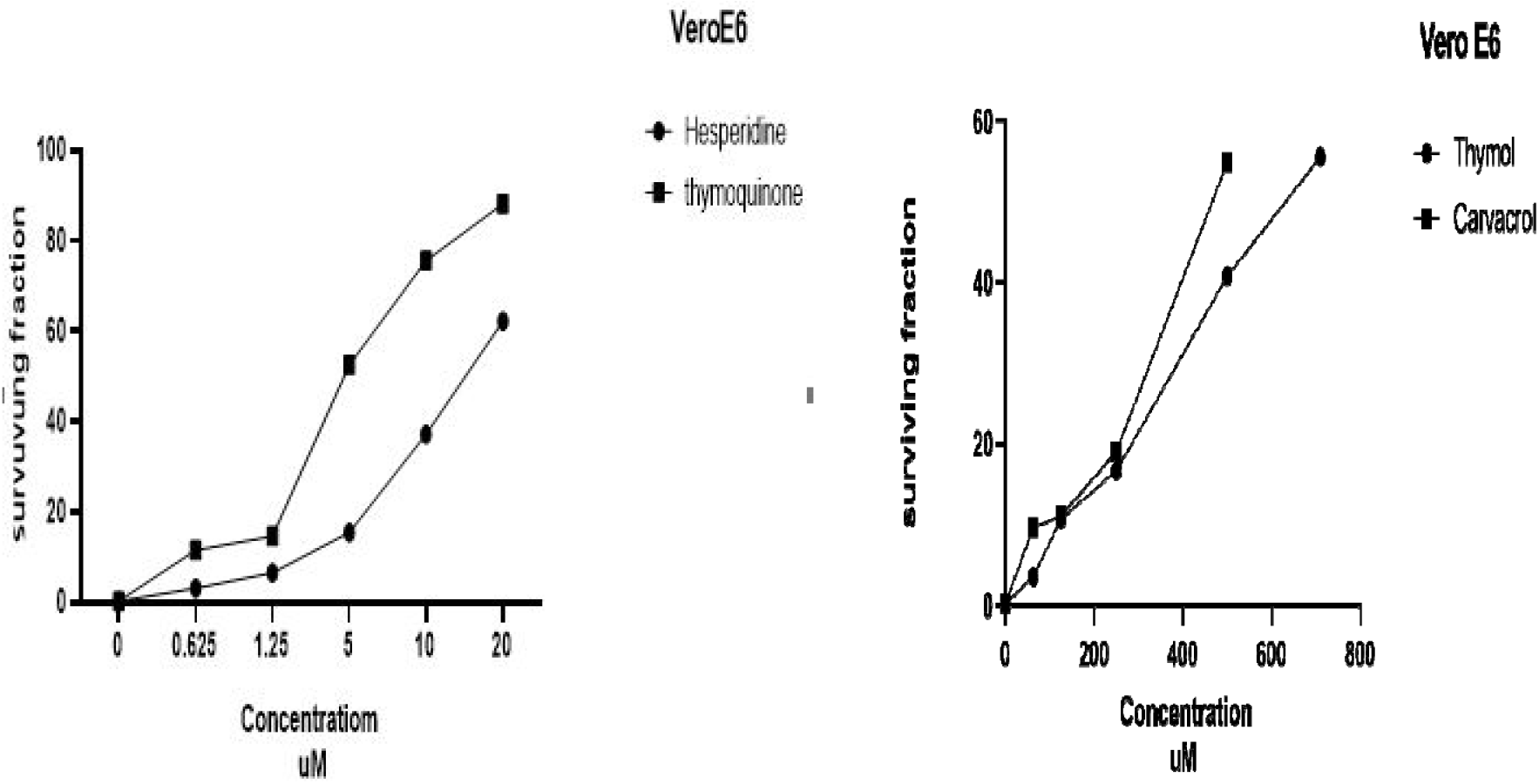
Effect of different concentrations Thymol, Carvacrol, Hesperidin and Thymoquinone on the cellular proliferation of Vero E6 cell line following 24 h of treatment. Values are expressed as the mean ± SD (n=3). Dose response curves were fitted using GraphPad prism 8.0.2 to determine IC50 values.

### In vitro study for accessing the antiviral effect of natural products against SARS-CoV-2

Our results revealed that Hesperidine was the most affective natural product against SARS-CoV-2 in Egyptians patients, as it caused inhibition to approximately 100% for infected Vero E6 cell line, Carvacrol, Thymoquinone caused inhibition to about 98%, Chloroquine as a positive control caused inhibition to 98% and Thymol show the lowest inhibition percentage 96% for infected Vero E6 cell line.

The results showed significantly difference between natural products effect with (P-value < 0.0001) as shown in figure 5.

**Figure 5:**
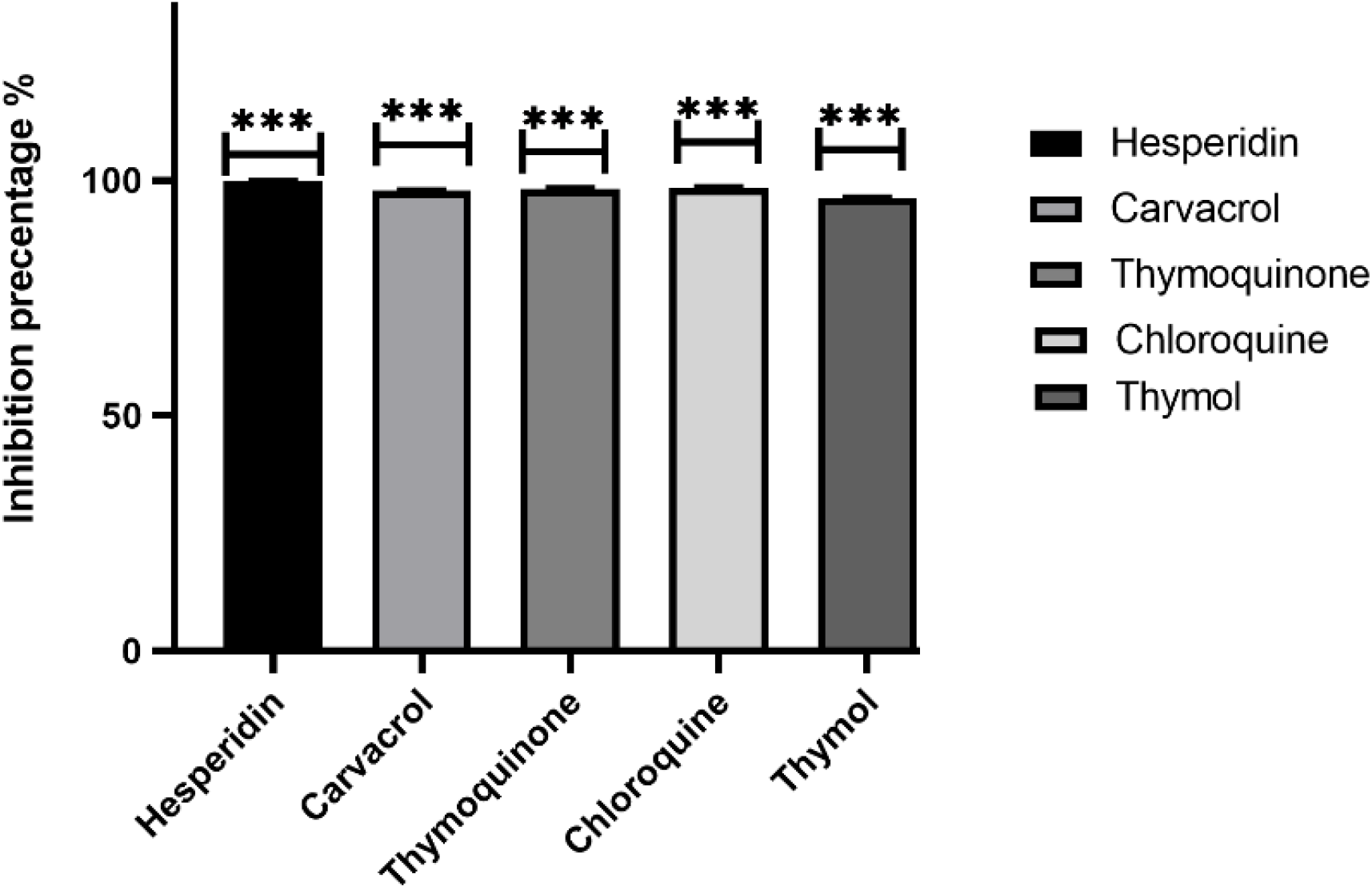
The antiviral effect of Hesperidin, Carvacrol, Thymoquinone, Chloroquine and Thymol against SARS-CoV-2 in vitro. The values are represented as means ± SD (n=2) (*p<0.0001).

These results indicate that Hesperidine and other natural compounds as Carvacrol and Thymoquinone could be therapeutic agent against SARS-CoV-2 in Egypt as treatment resulted in the effective loss of essentially all viral material by time.

Eventually, development of an effective anti-viral for SARS-CoV-2, if given to patients early in infection, could help to limit the viral load, prevent severe disease progression and limit person-person transmission. Benchmarking testing of those natural compounds against other potential antivirals for SARS-CoV-2 with alternative mechanisms of action would thus be important as soon as practicable.

## Competing Interest

All the authors declare that there is no competing interest in this work.

## Data Availability

The docking structures are available upon request from the corresponding author.

## Notes

### Competing Interest Statement

The authors have declared no competing interest.

